# Clustered Protocadherin *Cis*-interactions are Required for Homophilic Combinatorial Cell-Cell Recognition Underlying Neuronal Self-Avoidance

**DOI:** 10.1101/2023.11.11.566682

**Authors:** Gil Wiseglass, Nadir Boni, Karina Smorodinsky-Atias, Rotem Rubinstein

**Affiliations:** School of Neurobiology, Biochemistry and Biophysics, The George S. Wise Faculty of Life Sciences, Tel Aviv University, Tel Aviv, Israel; Sagol School of Neuroscience, Tel Aviv University, Tel Aviv, Israel

## Abstract

In the developing human brain, only 53 stochastically expressed clustered protocadherin (cPcdh) isoforms enable neurites from an individual neuron to recognize and self-avoid, while maintaining contact with neurites from other neurons. Cell assays have demonstrated that self-recognition occurs only when all cPcdh isoforms perfectly match across the cell boundary, with a single mismatch in the cPcdh expression profile interfering with recognition. It remains unclear however, how a single mismatched isoform between neighboring cells, is sufficient to block erroneous recognitions. In using systematic cell aggregation experiments we show that abolishing cPcdh interactions on the same membrane (*cis*) results in a complete loss of specific combinatorial binding between cells (*trans*). Our computer simulations demonstrate that the organization of cPcdh in linear array oligomers, composed of *cis* and *trans* interactions, enhances self-recognition by increasing the concentration and stability of cPcdh *trans* complexes between the homotypic membranes. Importantly, we show that the presence of mismatched isoforms between cells drastically diminishes the concentrations and stability of the *trans* complexes. Overall, we provide an explanation for the role of the cPcdh assembly arrangements in neuronal self/non-self-discrimination underlying neuronal self-avoidance.

## Introduction

A limited number of cell adhesion receptors play a major role in neuronal patterning during embryonic development. It seems unlikely that a relatively small group of proteins mediates such precise and complex processes, and indeed, it is becoming evident that these proteins rely on regulated expression and combinatorial interactions to carry out these processes (1–5). Among these adhesion receptors, the clustered protocadherins (cPcdhs) regulate neuronal survival (6), synaptogenesis (7–10), dendrite arborization (11), and neuronal tiling (12, 13). Clustered Pcdhs also mediate neuronal self-avoidance, a process that presents a unique challenge for cell-cell recognition and selectivity. Neuronal self-avoidance involves the recognition and avoidance of neurites originating from the same cell, while allowing neurites from different neurons to interdigitate and occupy the same field. This process requires neurons to distinguish “self” from “non-self” and is carried out by cPcdhs conferring unique cell surface identities to individual neurons.

In Mammals, approximately 60 *cPcdh* genes are organized in three tandem clusters: α, β, and γ (14, 15). Each neuron expresses a unique combination of isoforms on its cell surface, through stochastic promoter choice (14, 16–19). While all cPcdh isoforms share similar structures, they differ in their amino acids sequences and interact strictly homophilically in *trans* (i.e., between cells) (4, 20–25). Counterintuitively, this binding does not lead to adhesion, but rather triggers neurite-neurite repulsion. However, with anywhere between millions to billions of neurons within the mammalian brain and with fewer than 60 cPcdh isoforms, it is inevitable that many neighboring neurons will express several identical isoforms (26). Cell based assays showed that few mismatched isoforms that are not shared between neighboring neurons, could interfere with the homophilic recognition of identical isoforms, and thereby prevent incorrect repulsion. In these assays binding occur only between cells that express identical set of cPcdh isoforms, and a single mismatched isoform is sufficient to prevent cell-cell recognition (Figure 1). Thus, cPcdh isoforms can combine their specificities when co-expressed to determine cell-cell recognition.

**Figure 1.**
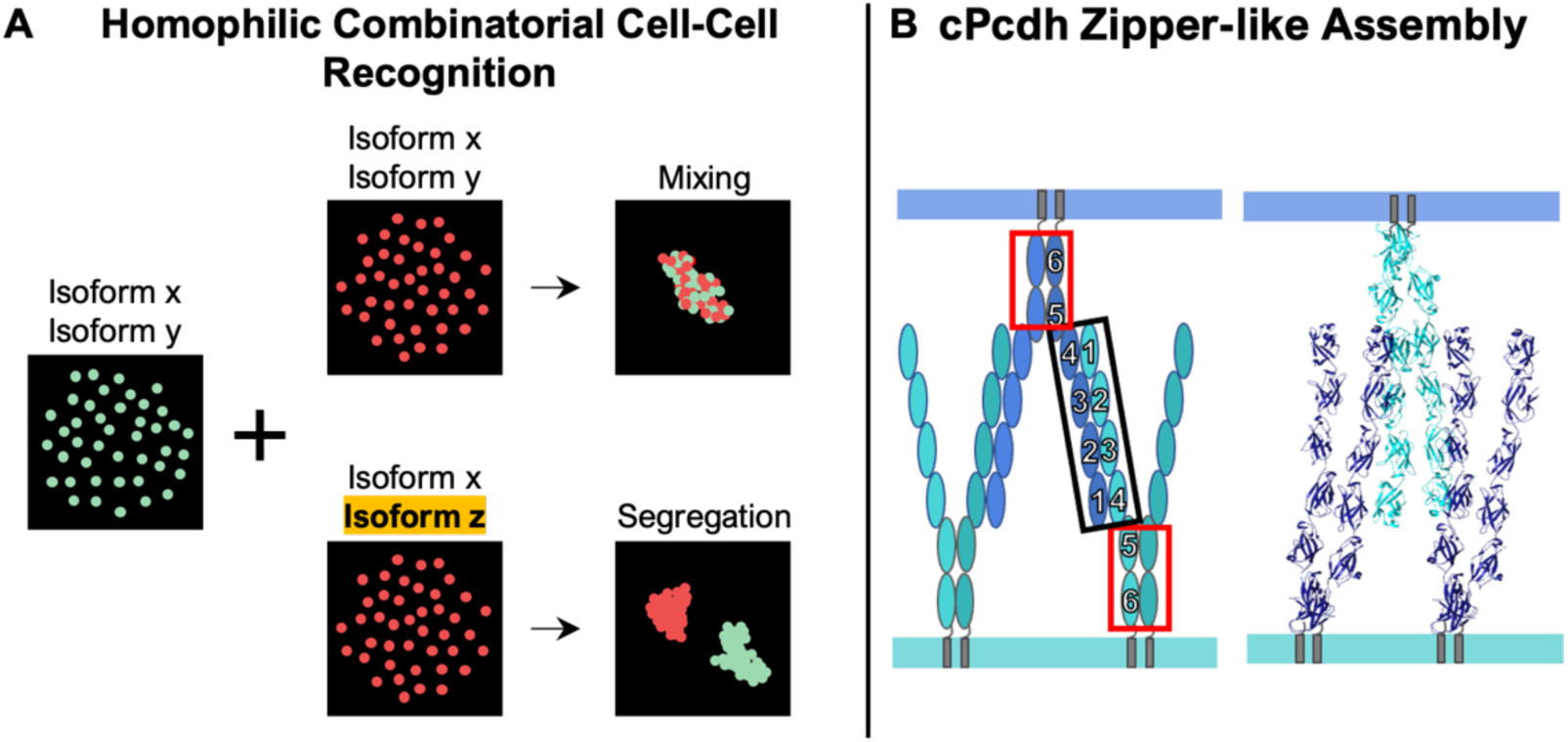
A) Schematic representation of the combinatorial homophilic binding in cell aggregation assays. Cells co-expressing a pair of distinct red fluorescently tagged cPcdh isoforms were assayed for interactions with cells co-expressing a pair of green fluorescently tagged isoforms. Mixed red and green coaggregates occur only between cells that express identical isoforms (Top). Separate green and red aggregates formed between cells expressing a mismatched isoform (Bottom). B) Illustration and the crystal structure of cPcdhgB4 cis dimer bound to two cis dimers of cPcdhgB4 from adjacent membrane (PDB code 6E6B (20). Cis interactions involving the membrane proximate EC5-EC6 domains (red rectangle) and *trans* interactions the EC1-EC4 domains (black rectangle). These arrangements result in a periodic zipper-like array that has been suggested to have a functional role in triggering the self-avoidance signal (29).

On the molecular level, structural and biophysical studies showed that in addition to strict homophilic *trans* interactions, cPcdhs engage in promiscuous *cis* interactions (on the same membrane) via an independent interface (4, 20, 27–29). Crystal structures and cryo-electron tomography (cryo-ET) of a single cPcdh isoform revealed the formation of zipper-like linear arrays at membrane contact sites formed by alternating *cis* and *trans* interactions (Figure 1) (29). These linear arrays align to create ordered two-dimensional superstructures that do not require any additional forces to operate between the zippers (29, 30). Although these zipper-like arrays are likely the recognition complexes formed between cells, previously they have only been determined at liposome contact sites, using a single isoform, and have yet to be linked to cell behavior. Thus, it has not been demonstrated whether these complexes can explain the combinatorial specificities observed in cells expressing multi-cPcdhs isoforms. Moreover, the effect mismatched isoforms have on the two-dimensional zipper-like cPcdh superstructures remains unknown.

In this study, we bridge this existing gap using an integrated approach that includes cell aggregation assays with multiple co-expressed wild-type or mutated isoforms and computer simulations. We show that *cis* interactions amplify cPcdh adhesion strength and are crucial for achieving combinatorial binding specificity. Using computer simulations, we demonstrate the significance of the zipper-like structure in modulating cPcdh concentrations at membrane contact sites. When all isoforms are identical, cPcdhs densely populate the contact between interacting membranes. In contrast, the presence of a mismatched isoform sharply reduces this concentration. Importantly, the concentration of cPcdh *trans* complexes, mediated by the zipper superstructures, is particularly responsive to the presence of mismatched isoforms. This enables cells to tolerate an increased expression of matching cPcdh proteins without incorrectly resulting in cell-cell binding. These findings provide a functional role for *cis* dimerization and the zipper-like superstructures in cPcdh combinatorial specificity.

## Results

### *Cis* interactions amplify cPcdhs *trans* adhesion

We utilized cell aggregation assays to experimentally test how cPcdh proteins combine their homophilic specificities. For these experiments we selected non-adherent K562 cells that lack the endogenous expression of cPcdhs and have already been used to study cPcdhs cell-cell recognition^4,20,27^. We selected eight cPcdh isoforms (α9, αC2, β5, γΑ1, γΒ2, γΒ5, γC3, and γC5) representing at least one member from each of the cPcdh clusters (α, β, γ) and sub-groups (γA, γB and C-type). To isolate the contribution of *cis* interactions to cPcdh combinatorial cell adhesion we introduced mutations that abrogate *cis* interactions. Previous studies have shown that in all isoforms cPcdh *cis* interactions involve the membrane-proximal EC5-EC6 domains (4, 27–29, 31). Importantly, deleting the EC6 domain only prevents the *cis* dimerization but not the *trans* dimerization, which is mediated by a different region of the protein (i.e., the membrane distal domains EC1 – EC4 (20)).

We initially tested whether deleting the EC6 domain, and thereby abrogating *cis* interactions, would also impact cell adhesion. We compared the behavior of K562 cells transfected with a single protein: wild-type cPcdh, ΔEC6 cPcdh mutant, or EGFP as a control. With the exception of the wild-type cPcdh-α9 that is unable to reach the cell surface when expressed alone (4, 32, 33), cells expressing either the wild-type or the ΔEC6 ectodomains aggregated for all the tested isoforms. In contrast, the control cells expressing EGFP did not form aggregates (Figure 2). These findings were consistent with our expectations, based on previous results, that abolishing cPcdh *cis* interactions would not prevent *trans* interactions (4, 20). Interestingly, here we observed that cells expressing ΔEC6 cPcdh generally formed smaller and fewer aggregates compared to the aggregates formed by their wild-type counterparts (Figure 2). The range of cumulative size of the aggregates generated by the ΔEC6 mutants was between 9-50% of their wild-type counterparts. Of the eight isoforms, only cPcdhγB2 formed similar sized aggregates for both the wild-type and the mutant constructs (Figure 2). It is possible that for this particular isoform, a combination of high expression levels and a strong *trans* binding affinity compensated for the lack of *cis* interactions caused by the deletion of the EC6 domain. However, these findings overall suggest that *cis* interactions increase the adhesive strength of cPcdhs.

**Figure 2.**
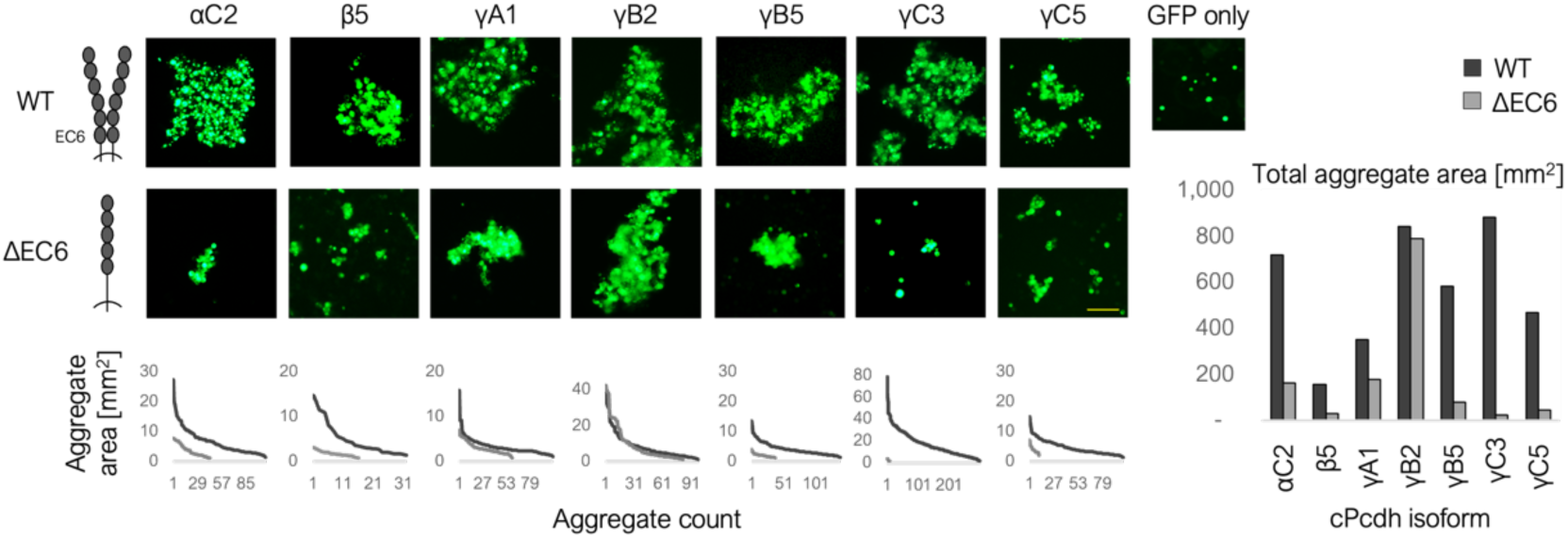
Abrogating the *cis* interactions results in reduced size of aggregates. Representative results of aggregations of cells transfected with a single cPcdh isoform, either wild-type or ΔEC6 ectodomains, or GFP are shown. Both specific and total aggregate sizes for each isoform are shown in the bottom row and as a bar plot on the right, respectively. Scale: 100 µm.

### *Cis* interactions are essential for cPcdh combined adhesion specificity

Previous studies used cell aggregation assays to show that cPcdhs combine their homophilic adhesion specificity to generate a new cell surface identity (4, 27) (Figure 1). Here we tested the effect of cPcdh *cis* interactions on this combinatorial specificity by using aggregation assays with mismatched wild-type or mutated isoforms. Specifically, we tested the binding preferences for cells co-expressing two isoforms fused to EGFP with cells co-expressing a common isoform and a mismatched isoform fused to mCherry. Figure 3A shows the cell aggregation images, with the top images corresponding to cells co-expressing two wild-type isoforms capable of both *cis* and *trans* interactions. The bottom images correspond to cells co-expressing ΔEC6 isoforms unable to form *cis* interactions. To quantify the degree of aggregate mixing, we utilized a custom Python script calculating the proportion of neighboring red and green cells (see Methods, and Figure S1). This proportion (referred to as the mixing score) is shown in the corner of each image, with a value greater than 0.1 indicating visibly mixed aggregates (Figure 3 and Figure S1).

**Figure 3.**
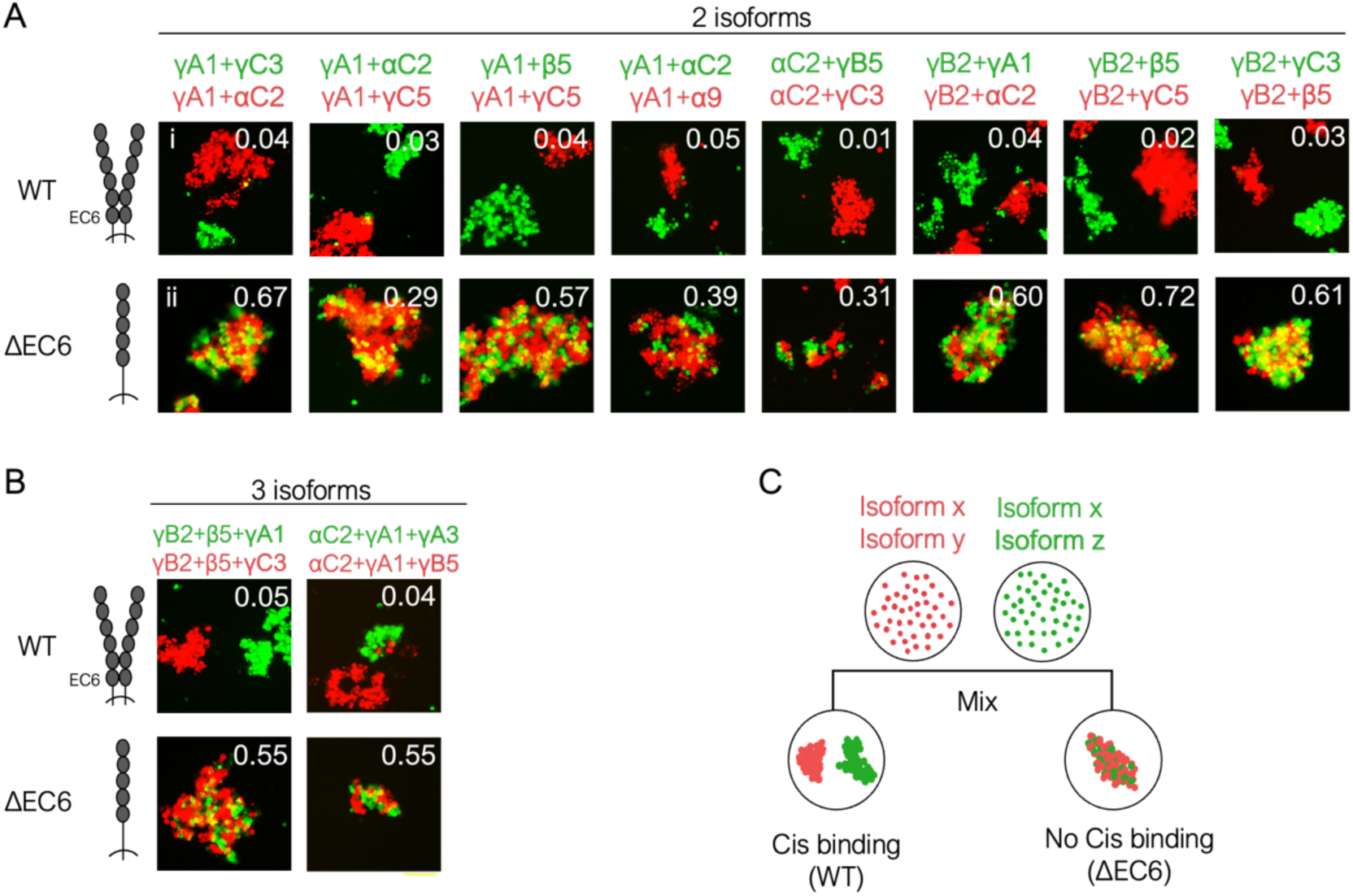
Mismatch coaggregation assays demonstrate the essential role of *cis* interactions in cell-cell combinatorial homophilic binding specificity. (A, B) Red and green cells co-expressing matched and a mismatched wild-type cPcdhs did not bind to each other and formed separate homotypic aggregates (top). In contrast, eliminating *cis* binding leads to indiscriminate adhesion and mixed coaggregates between red and green cells expressing mismatched isoforms (bottom). The mean mixing index values are shown in the upper right corner of each representative image (see also Figure S1). (C) Illustration of the cell aggregation experiment. Scale: 100 µm.

In Figure 3 each column represents a different set of isoform combinations. For example, the first column shows a green cell population co-expressing cPcdh-γA1 with cPcdh-γC3 mixed with a red cell population co-expressing cPcdh-γA1 (a common isoform) with cPcdh-αC2 (a mismatched isoform). With the wild-type isoforms, the two cell populations did not mix, instead they formed separate red and green aggregates with a mixing index of 0.04 (Figure 3Ai). These findings correspond to previous observations of cPcdhs combinatorial binding specificity where the presence of a mismatched isoform prevents coaggregation^4^. Remarkably, when we carried out cell adhesion experiments using the same isoform combinations, with the ΔEC6 mutation, the two cell populations produced mixed red and green aggregates with mixing index of 0.67 (Figure 3Aii). We observed this behavior, where the *cis* interaction is essential for combinatorial binding specificity, with different isoform combinations (Figure 3A). Specifically, cells expressing identical sets of multiple ΔEC6 isoforms formed mixed red and green aggregates (Figure S2). Thereby, demonstrating that eliminating *cis* interactions does not abrogate *trans* recognition, when multiple identical isoforms are expressed, but rather impacts the ability of cPcdhs to combine different *trans* specificities.

We also assessed the impact of cPcdh *cis* interactions on cell-cell recognition for cells co-expressing three isoforms. Cells expressing wild-type cPcdhs with a mismatched isoform formed separate red and green aggregations (Figure 3B top). In contrast, cells expressing ΔEC6 cPcdh constructs formed mixed red and green aggregates even in the presence of mismatched isoforms (Figure 3B bottom). Thus, overall cells expressing mismatched cPcdhs that are unable to interact in *cis*, form heterotypic (mixed) aggregates, which is a strikingly different behavior compared to cells expressing wild-type cPcdhs that form separate homotypic aggregates (Figure 3C). These results point to the critical role of cPcdh *cis* interactions in combinatorial cell-cell recognition specificity.

### Computer simulations reveal the connection between *cis* interactions, zipper-like assemblies, adhesion strength and combinatorial selectivity

We developed a lattice-based computer simulation of cPcdh interactions in the adhesion contact of two cell membranes, aimed to elucidate how the zipper-like assembly impacts cell adhesion strength and selectivity (Figure 4A). We began by simulating cells expressing a single isoform. In the cell aggregation assays described above (Figure 2), cells expressing a wild-type isoform formed significantly larger aggregates compared to cells expressing an isoform unable to interact in *cis*. To gain insight into this behavior, we utilized our simulation method to computationally replicate these assays. Consistent with our previous report, the presence of both *cis* and *trans* interactions resulted in the formation of long zipper-like assemblies aligned in two-dimensional arrays. In contrast, in the absence of *cis* interactions, *trans* dimers continued to form, but without ordered oligomers (Figure 4B) (30). Notably, the concentration of *trans* dimers decreased significantly in the absence of *cis* interactions (Figure 4 B & C). In cell aggregation assays a reduction in the number of *trans* dimers results in a weaker adhesion between cells and therefore smaller aggregates.

**Figure 4.**
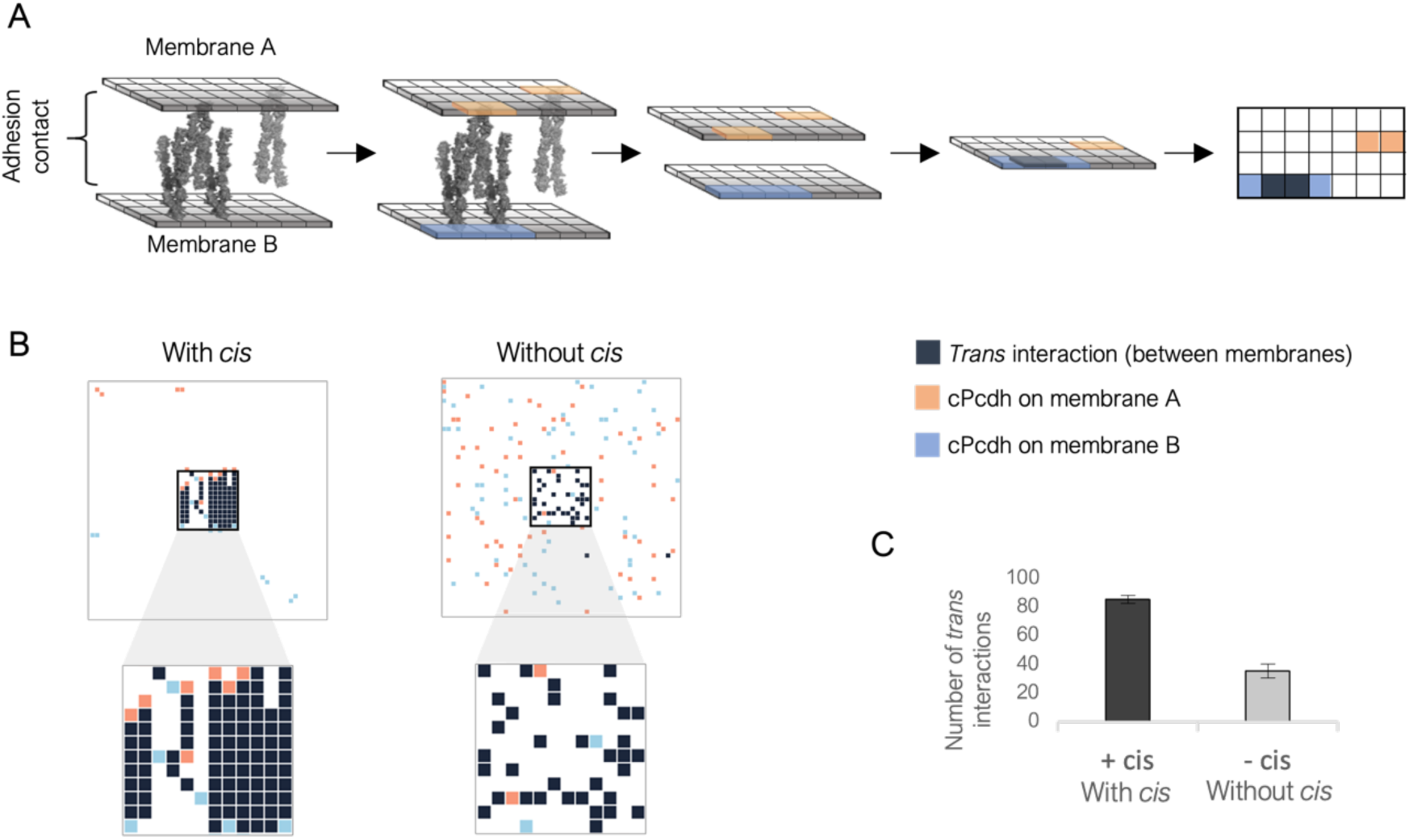
Computer simulations reveal how *cis* interactions and the zipper-like assemblies impact adhesion strength. A) Illustrate the lattice simulations, designed as two stacked grids representing both membranes. Squares on the grid occupied by cPcdhs are colored based on membrane affiliation, with orange representing membrane A and blue representing membrane B. *trans* dimers are indicated by black squares (i.e., zipper-like arrays are represented as black lines). B) Simulation snapshots of the 2D lattice model when both *cis* and *trans* interactions are present (left) and when *cis* interaction is absent (right). *Trans* interactions can only occur in the diffusion trap area shown in the middle and magnified. Only with *cis* interactions, 2D stacking of zipper-like structures are present at the diffusion trap area. C) Boxplot of 60 independent simulations with or in absence of *cis* interactions demonstrate a reduction in *trans* dimers when *cis* interactions are abolished.

To clarify the impact of *cis* interactions on combinatorial homophilic interactions observed in mismatched cell aggregation assays (Figure 3), we computationally modeled the interactions between cells expressing multiple distinct isoforms. Our simulation mimicked the two types of competing cell-cell interactions, occurring in the mismatched aggregation assay: The first involves adhesion between homotypic cells, such as the green cells adhering to other green cells. In this scenario, these cells express an identical pair of two isoforms. The second type involves adhesion between heterotypic cells (i.e., interactions between red and green cells) expressing a combination of an identical isoform and a mismatched isoform.

Our simulations of wild-type isoforms revealed that homotypic binding (i.e., cells expressing identical isoforms) results in densely populated *trans* complexes that are organized in two-dimensional long zippers-like arrays (Figure 5Ai). In contrast, a mismatched isoform between heterotypic cells results in the formation of shorter, one-dimensional zipper-like assemblies (Figure 5Aii) and significantly fewer *trans* complexes (Figure 5C top). Notably, in the absence of *cis* interactions, the introduction of a mismatched isoform did not change the cPcdh assemblies, as zipper-like arrays were absent and there was only a slight reduction in the number of *trans* complexes (Figure 5B i & ii, 5C top).

**Figure 5.**
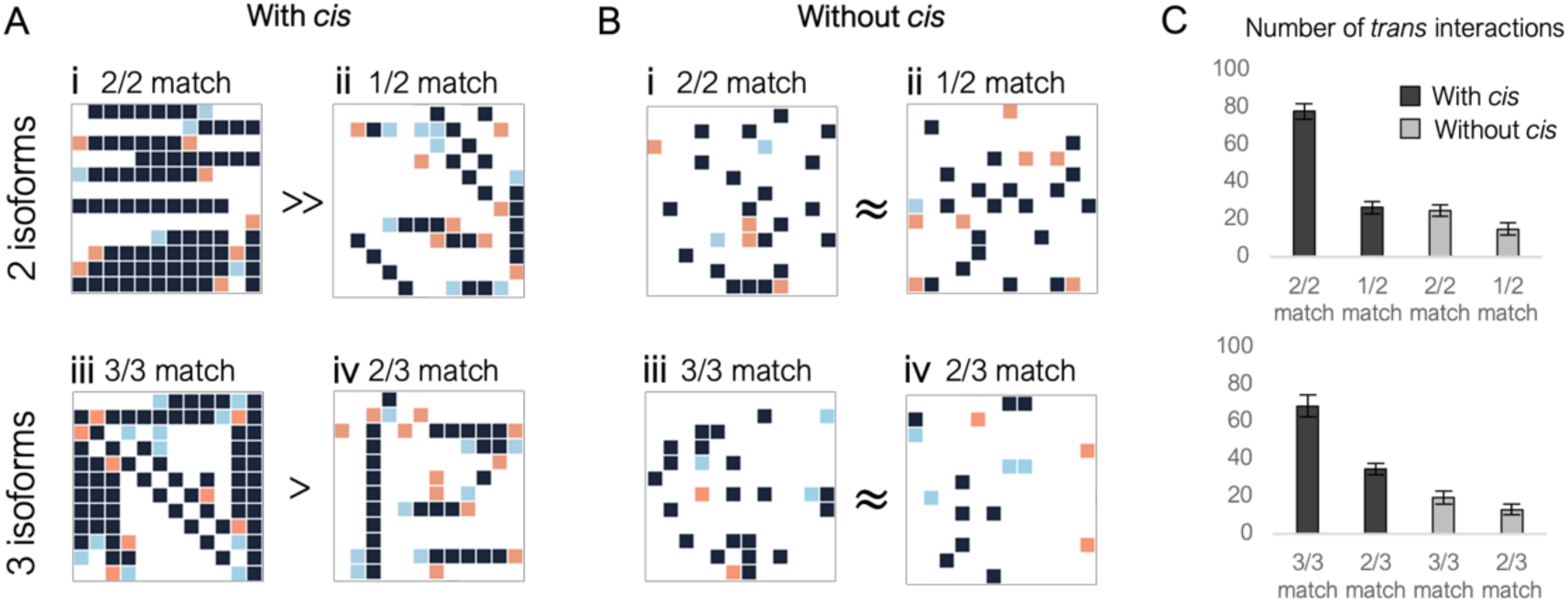
Computer simulations reveal the role of *cis* interactions and the zipper-like assemblies in adhesion combinatorial specificity. Snapshots of simulations for interactions of wild-type (with *cis*, A) and ΔEC6 (without *cis*, B) cPcdhs. Top images represent the results of simulations with two isoforms (i & ii), and bottom images represent results of simulations with three isoforms (iii & iv). Only the diffusion trap areas are shown. (A) Simulation of wild-type matching cPcdhs generated clusters of long zippers-like arrays and significantly more *trans* dimers compared to simulations of cells that expressed a mismatched isoform. (B) Once *cis* interactions are eliminated, a mismatched isoform only slightly reduces the number of *trans* dimers compared to simulations of all matched isoforms. C) Boxplots demonstrating the number of *trans* dimers observed from simulations of different match to mismatch ratios with and without *cis* interactions (30 independent simulation run for each condition).

To further characterize the impact of *cis* interactions on adhesion selectivity in mismatched aggregation assays, we simulated cells expressing three isoforms mimicking the cell aggregation assays described in Figure 3B. We observed similar results, specifically with *cis* interactions the concentrations of cPcdh *trans* complexes significantly diminish in the presence of mismatched isoforms (Figure 5A iii & iv, 5C bottom). In contrast, in the absence of *cis* interactions, the concentration of *trans* complexes is similar for bindings between homotypic and heterotypic cells. (Figure 5B iii & iv, 5C bottom).

### *Cis* interactions resist incorrect cell recognition also at increased expression of matched isoforms

Aggregation assays of cells expressing multiple cPcdh isoforms have shown that a mismatched isoform prevents coaggregation (see Figure 3 and references (4, 27)). However, increasing the expression of the matched isoforms results in the formation of mixed aggregates, suggesting that cPcdh recognition is affected by isoform identities and their expression level (27). These results align with prior studies of adhesion proteins linking expression levels and adhesion strength (2, 34). We used our simulations to quantify the ability of cells to tolerate increased expression of the matched isoform without wrongly binding to cells that also express mismatched isoforms. In addition, we tested the impact of *cis* interactions on this tolerance.

We performed eleven sets of simulations, modeling protein-protein interactions between cells expressing different ratios of matched-to-mismatched isoforms (0:10, 1:9, 2:8, …, 9:1, 10:0, respectively). Initially we only expressed the mismatched isoform and gradually altered the expression ratio in favor of the matched isoforms, similar to solution titration. We repeated each set of simulations to model protein-protein interactions for the following conditions: heterotypic cell-cell interactions (both cells express a matched isoform and a mismatched isoform), homotypic cell-cell interactions (both cells express the same set of isoforms), wild-type cPcdhs, and mutant cPcdhs unable to interact in *cis*. We used the results from these simulations to predict experimental cell aggregation patterns by comparing the number of *trans* dimers of homotypic to heterotypic cell-cell interactions. We reasoned that when the number of *trans* dimers between homotypic cells is similar to that between heterotypic cells, all cells will adhere in similar strength. Consequently, red and green mixed aggregates will form. In contrast, when there is a considerably higher number of *trans* dimers between homotypic cells compared to that between heterotypic cells, homotypic cells will adhere more strongly. This scenario would result in the formation of separate, homotypic aggregates (all-red aggregates and all-green aggregates).

In the case of the wild-type cPcdhs, we observed that until the expression of the matched isoform reached 70% of the total cPcdhs, there were significantly more *trans* dimers between homotypic cells compared to heterotypic cells (Figure 6B, black curve & Figure S3). Once the matched isoform constituted more than 70% of the total cPcdhs, the number of *trans* dimers between heterotypic cells dramatically increased and became similar to the number observed for homotypic cell bindings (Figure 6B, black curve). We used these results to predict that in cell aggregation assays, mixed aggregates would form only when the wild-type matched isoforms constitute more than 70% of the total cPcdhs. In contrast, when simulating cells expressing isoforms that do not interact in *cis*, we observed that the number of *trans* dimers between homotypic and heterotypic cells became similar when the matched isoform consisted of at least 40% of the total proteins (Figure 6B, gray curve). We expected that cells would form mixed aggregates for this value, which corresponds to half of the value observed in simulations of wild-type cPcdhs. Notably, for simulations of wild-type proteins, when *cis* dimerization occurs, the slope of the curve resembled a step function, suggesting a binary threshold for tolerating the presence of matching isoforms in heterotypic interactions (Figure 6B, black curve).

**Figure 6.**
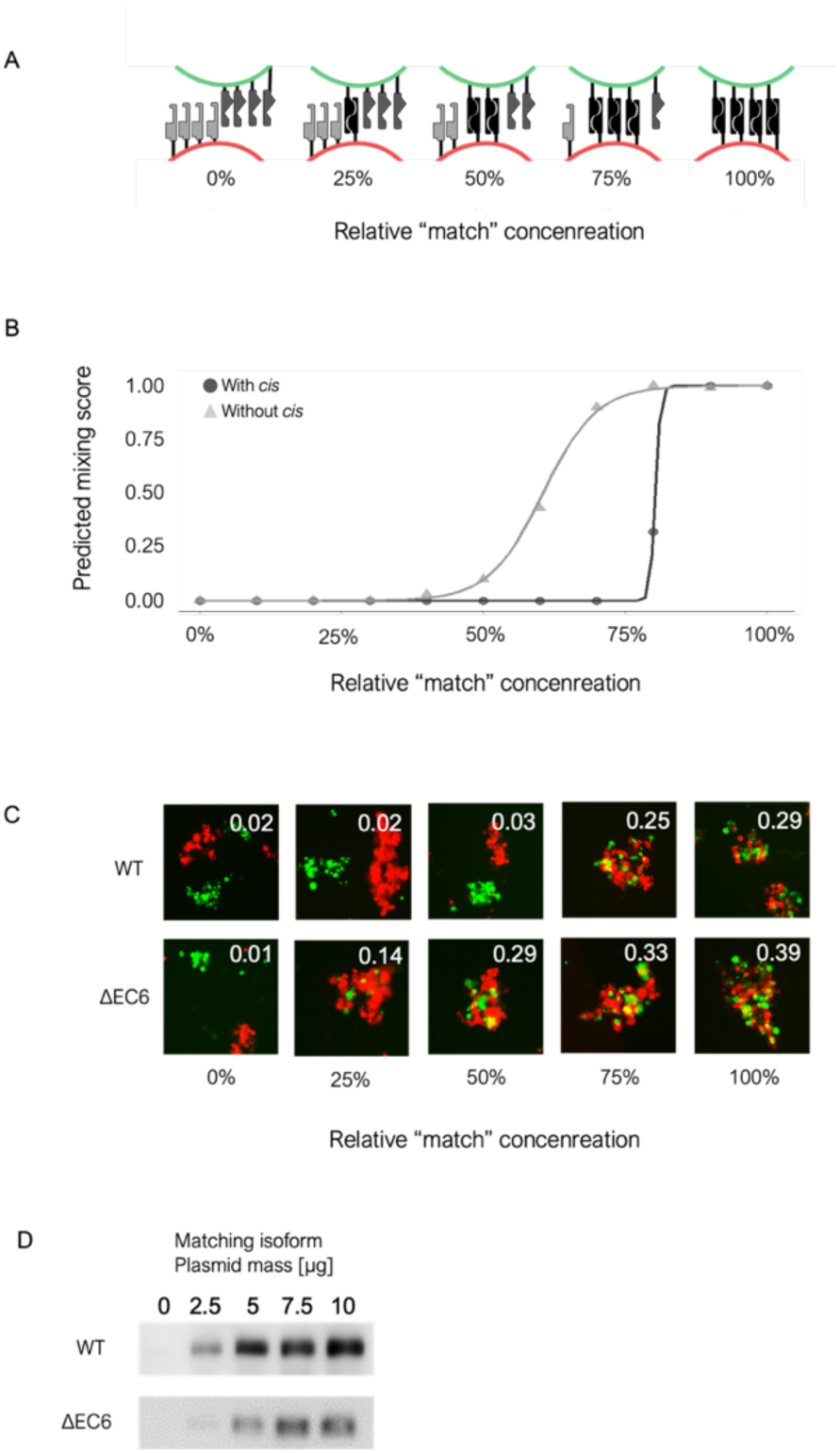
*Cis* interactions endowed tolerance to increased level of expression of matching isoforms. A) Illustration of the changing ratios of the matched to mismatched isoforms under five experimental conditions. Starting from the left, two distinct cell populations (represented in red and green) express two different isoforms, resulting in a 0% match. Next, both populations express two isoforms each, with one matching isoform. The concentration of the matching isoforms relative to all isoforms gradually increases from 25% to 100%. B) A summary of eleven sets of simulations, modeling the protein-protein interactions between cells with different ratios of matching isoforms. As the concentration of the matching isoform increases, the extent of mixing is predicted to increase gradually when *cis* interaction is absent (light grey). In contrast, with *cis* interactions, the slope of the curve resembles a step function, suggesting a binary threshold for tolerating the presence of matching isoforms (black). C) Cells transfected with increasing levels of the matching isoforms were assayed for co-aggregation. D) Western blot analyses of increased matching isoform expression, correlated to the amount of cPcdh plasmid DNA transfected (see also in Figure S3).

Remarkably, cell aggregation assays with mismatched isoforms recapitulated the simulation results. We experimentally tested five ratios of common to mismatched isoforms (0:4, 1:3, 2:2, 3:1, and 4:0, respectively; Figure 6A). We altered the ratio of common isoforms by adjusting the amount of plasmid DNA for each cPcdh construct used in transfection. This method has been well established (2, 27), and we validated transfection levels using Western blot analysis (Figure 6D & Figure S3). We found that wild-type cPcdhs expressing cells tolerated the increasing expression of matched isoform significantly better than the mutated cPcdhs that were lacking *cis* interactions (Figure 6C). Cells expressing wild-type matched and mismatched cPcdhs bound to each other and formed mix aggregates only when the matched isoforms were at least 75% of the total transfected cPcdhs (ratio of 3 matched to 1 mismatched isoform; Figure 6C top). In contrast, cells expressing mutated isoforms formed mixed aggregates already at the low ratio of 25% (1 matched to 2 mismatched isoforms; Figure 6C bottom). Notably, in the case of wild-type isoforms mixing of red and green cells occurs abruptly and in accordance with the step function predicted by the simulations. In contrast, without *cis* interactions, mixing of red and green cells occur gradually with the increased expression of matched isoforms. In summary, by quantifying the number of *trans* interactions observed in our simulations, we explained and predicted the experimental cell aggregation results.

## Discussion

This study examined the molecular basis of cPcdhs combinatorial cell-cell recognition. We focused on understanding how cells expressing a matched set of cPcdh isoforms selectively adhere to each other but not to cells expressing even a single mismatched isoform. This behavior is crucial for neuronal self-avoidance. Our work consisted of computational simulations and cell aggregation experiments and revealed several key insights: First, cPcdh *cis* interactions significantly amplify adhesion strength. Second, *cis* interactions are essential for cPcdh homophilic combinatorial binding. And third, *cis* interactions enable cells to tolerate an increased expression of matching cPcdh proteins without incorrect cell-cell binding. As will be discussed below, our simulations suggest that all these behaviors are driven by the zipper-like arrangements that lead to a dense *trans* complexes population at membrane contact sites only when all isoforms are identical.

Clustered Pcdh homophilic *trans* interactions promote cell binding via the EC1-EC4 domains (4, 20), while homophilic and heterophilic *cis* interactions that contribute to the formation of cPcdh zipper-like assemblies involve the EC5-EC6 domains (4, 23, 28, 29). These zipper-like assemblies are likely responsible for cPcdh-mediated combinatorial homophilic cell-cell recognition. Albeit, establishing this connection has proven challenging due to the complexity of a system involving multiple distinct adhesion proteins that interact in both *cis* and *trans*. To tackle this challenge, we isolated the contribution of the *cis* interaction to combinatorial cell-cell recognition by analyzing the aggregation of cells expressing wild-type or mutated cPcdh proteins that cannot bind in *cis*. While we experimentally tested eight representative isoforms, we note that based on previous sequence conservation, structural, and biophysical analyses most cPcdh proteins have similar features and binding properties. Therefore, our findings likely represent a generalizable phenomenon. Additionally, we developed a computer simulation that models the formation of cPcdh zipper-like arrays at the contact area between two cells. While the interactions between cPcdhs have three-dimensional features, we used a two-dimensional lattice model to reduce the number of parameters and computational complexity. Nevertheless, this seemingly straightforward model captured non-trivial experimental observations and explained the role of *cis* interactions and zipper-like assemblies on adhesion combinatorial selectivity.

In the less complex case of a single isoform expression, we found that preventing *cis* interactions lead to a decrease in aggregate size, indicating reduced adhesion strength. Notably, when we computationally simulated this scenario, we observed significantly fewer *trans* interactions between proteins lacking *cis* binding. The formation of relatively small aggregates can be understood when considering that the number of *trans* interactions correlates with adhesion strength, which in turn affects aggregation size. We note that previous studies report on the impact of *cis* interactions on cPcdh cell adhesion, however these reports only focus on *cis* interactions involving isoforms from the alpha cluster (4, 32, 33). The impact of this *cis* interaction on *trans* adhesion is trivial as alpha isoforms cannot reach the cell surface when expressed alone.

In the more complex scenario involving the expression of multiple isoforms per cell, we showed that eliminating *cis* binding leads to indiscriminate adhesion and mixed aggregates, even in the presence of a mismatched isoform. This is in stark contrast to the behavior of wild-type cPcdhs that only form mixed aggregates when all isoforms are matched. Using computer simulations of wild-type cPcdhs, we demonstrated that *cis* interactions and zipper-like arrangements significantly increase the concentrations of cPcdh complexes between cells expressing the same cPcdh isoforms. Notably, introducing even a single non-matching isoform significantly lowers the concentrations of these complexes and shortens the zipper-like chains. Remarkably, once the *cis* interactions are eliminated, the presence of a mismatched isoform only slightly reduced the number of *trans* interactions.

These findings can be understood using the Differential Adhesion Hypothesis (DAH), which proposes that the strength and specificity of cell-cell binding are determined by contacts that maximize the number of interactions across cell membranes (34, 35). According to the DAH, if the adhesion between homotypic cells is considerably stronger than that between the heterotypic cells, the cell population will form separate homotypic aggregates. This, in fact, was our observation with cells expressing mismatched wild-type cPcdhs. In contrast, when the homotypic adhesion is similar or marginally stronger than the heterotypic adhesion, the cells will bind to each other and aggregate via both homotypic and heterotypic interactions. This was also consistent with our observations of cells expressing a mismatched isoforms lacking *cis* dimerization.

### Relevance to *in-vivo*

In our computer simulations, cPcdh formed two-dimensional clusters by alignment of zipper-like arrays only when all isoforms were matched. These clusters likely enhance the *trans* complexes’ stability, which could explain the increase in the number of *trans* interactions, and consequently, the stronger adhesion between homotypic membranes. Clustering, as a means of stabilization of *trans* complexes, has been described for both classical (34, 36, 37) and atypical cadherins (38).

Thus, our findings explain how the zipper-like structures could provide neurons with the necessary cell surface diversity for self-avoidance. *In-vivo*, these large clusters of cPcdh complexes would presumably form temporarily and deconstruct, as they are only generated between sister neurites to trigger repulsion. Similar to other adhesion proteins, mechanisms including protease cleavage of cPcdhs (32, 39–41) or trans-endocytosis (38, 42–44) could be involved in remodeling membrane boundaries by deconstructing stable cPcdh complexes.

Another complexity of cPcdh function *in-vivo* is the considerable variability in expression levels of different isoforms. For example, Purkinje neurons have a constitutive and higher expression of C-type cPcdhs. This expression pattern could result in neighboring neurons with high expression levels of identical isoforms that, in turn, could impact the appropriate discrimination between self and non-self interactions. However, a recent report suggests that a bias for preferential *cis* dimerization between cPcdhs from different subclusters could reduce potential misrecognition by increasing cell surface diversity (31). Our current findings also contribute to an alleviation of this issue. We demonstrated that the formation of zipper-like cPcdh structures at the cell-cell contact area provides cells with resistance to cell-cell misrecognition also in the presence of highly expressed matched isoforms (Figure 6). When we removed *cis* interactions that prevent the formation of zipper-like structures, cells transfected mostly with mismatched isoforms (common isoform were only 25% of the expressed cPcdhs) were unable to tolerate the presence of the common isoform and bound to each other. We note that the low tolerance (less than 25%) measured for cPcdhs without *cis* interactions is similar to the estimated tolerance of the Fly Dscam1 (10 – 20%) that behaves as monomer and mediates neuronal self-avoidance in the fly (45).

Future research should address further complications within the cPcdh system. For instance, interactions between cPcdh and other proteins, both extracellularly (e.g., RET and Neuroligin (46, 47)) and intercellularly (e.g., Axin1 and PYK2 (48, 49)). These interactions likely contribute to the diverse neuronal patterning functions mediated by cPcdhs. In addition, previous studies have shown that each neuron can express up to 15 different cPcdh isoforms (50). However, the critical question of how many of these isoforms must match between two neurons to trigger a misrecognition as ‘self’ remains unclear. This is mainly due to the technical difficulty of co-expressing more than a handful of isoforms in experimental settings. Under this limitation, our cell aggregation results validate the accuracy of the computer simulation model in predicting selective cell-cell recognition. Consequently, similar simulation methods that incorporate a larger number of cPcdh isoforms could provide valuable insights into the threshold for misrecognition and the complexity of self-recognition mechanisms mediated by cPcdhs.

While our study focused on cPcdhs, many other adhesion receptors utilize a combination of *cis* and *trans* interactions to form zipper-like arrays at the cell-cell boundaries (51). Our experimental and computational setup could be extended to understand how these adhesion proteins induce cell-cell recognition and patterning.

## Materials and Methods

### Plasmid construction

As with previous studies, we found that cPcdh constructs that included the extracellular and transmembrane regions but were missing most of the cytoplasmic region, expressed better and exhibited bigger consistent aggregates compared to full-length proteins (2, 27). Therefore, all the experiments in this study included cPcdhs with a truncated cytoplasmic region. Constructs used for the cell aggregation experiments in this study were kindly contributed by Prof. Joshua A. Weiner from Iowa Neuroscience Institute, department of biology, the University of Iowa (cPcdhγA3, γB2, γC3 and γC5). The remaining isoforms were purchased from DNASU (MmCD00083158, MmCD00295450, MmCD00295385, MmCD00295452 and MmCD00295264). The coding region of each cPcdh isoform was inserted into a pcDNA3.1(+) with either EGFP or mCherry in the 3’ end, without most of the cytoplasmic domain. For deletion of the EC6 domain, the entire constructs except for the EC6 domains were amplified using EcoRV restriction sites in the 3’ of the primers. The PCR products were than digested with EcoRV and ligated using T4 DNA ligase. For expression analysis by Western blot, HA tags were added to the C-terminus of EGFP or mCherry of the cPcdh γB2 constructs, using Gibson assembly.

### Cell aggregation assay

K562 cells were cultured in IMDM medium supplemented with 10% FBS, GlutaMAX™ and Pen-Strep (0.02 mg/ml Streptomycin sulfate and 20 units/ml Penicillin G sodium salt). For each transfection reaction, one million cells were utilized and transfected with a total of 10 µg plasmid DNA, using the Amaxa Nucleofector II (K562 ATCC program). The electroporation protocol followed a similar procedure to Chicaybam et al., 2013 (52), using buffer “1M”. In the single isoform experiments (Figure 2), cells were treated with 3 mM calcium chloride (at a final concentration) 1-3 hours post transfection. The morning following transfection, 500 µl of cells were transferred and mixed in 6-well plates with 1.5 ml of complete DMEM supplemented as mentioned earlier, along with 3 mM calcium chloride. For the isoform mismatch and different isoform ratio experiments (Figures 3, 6 and S2), 250 µl of cells from two different transfection wells (totaling 500 µl) were combined in 6-well plates with 1.5 ml of complete DMEM, supplemented as described earlier, and including 3 mM calcium chloride. In the isoform mismatch experiment, the same plasmid ratio was maintained between the wild-type and ΔEC6 combinations. In this experiment, cells were additionally supplemented with 4 mM valproic acid. Valproic acid, a compound known for enhancing recombinant protein expression in cell line cultures, was utilized for this purpose without effecting cell adhesion (2, 52–56). After mixing, the cells were incubated (37°C, 5% CO_2_) on a rocking platform at 15 RPM for 2-4 hours. Images were acquired using Nikon Eclipse Ts2 and Eclipse Ti inverted microscopes.

### Results quantification

Aggregate size for the results presented in Figure 2 was measured using ImageJ^56^ freehand selection tool in 64 images per construct. Aggregate mixing or separation presented in Figures 3 and 4 was quantified using a custom Python script. This script calculates the ratio of red and green cells that are in close proximity to each other. To achieve this, the script divides the red and green channel images into a grid and examines corresponding grid cells to identify the presence of both a red and a green K562 cell. Each grid cell’s size is equivalent to ∼1.5 K562 cells (24.32 µm²). Therefore, if both the red and green K562 cells occupy the same grid cell, it suggests potential contact between them. The proportion displayed for each image represents the average score obtained from a minimum of two biological experiment replications. Within each replication, 4-6 images were captured and analyzed. For additional statistical details, see supplemental figure 1.

### Western blot

Nitrocellulose membranes were blotted with Anti-HA-Tag Rabbit Monoclonal Antibody (Cell Signaling #3724) and Anti-rabbit IgG, HRP-linked Antibody (Cell Signaling #7074). Signal emitted by protein bands was quantified using ImageJ (57) gel lane-plotting tool. Bar chart summary of three replications for each lysate sample, normalized for total-protein quantification by ponceau staining.

### Computer simulations

In our simulation, the cell-cell contact is represented by two stacked grids (size of 50×50), with cPcdhs randomly assigned to them during the initiation step. The system then reaches equilibrium by using the Monte Carlo method. In each Monte Carlo step, the simulation can perform one of six actions (translational or rotational movement, association or disassociation in both *cis* and *trans*) randomly. To visualize the equilibrium state, the cPcdhs are presented as squares with colors indicating membrane affiliation (orange for membrane A, blue for membrane B, and black for a location with double occupancy from both membranes). Figure 4A briefly demonstrates the simulation design and visualization process, and for a more detailed description see our previous work (30). All the simulations were performed under a total cPcdh concentration of 4% of the total grid area. The contact area was simulated by a squared diffusion trap in the center of the grid with an area of 5% of the total grid size. The *cis* and *trans* affinities were ΔGD(*cis*) = 7kT and ΔGD(*trans*) = 5kT. The source codes are available for download at https://github.com/Rubinstein-Lab.

### Prediction of mixing score

We used our simulation model to predict the mixing score between heterotypic cells as determined by cell aggregation assay results. As the number of interactions is a proxy for adhesion strength, we compared the contacts of homotypic (full isoform match) and heterotypic (a single isoform mismatch) cell-cell interactions. We performed multiple independent simulation runs for each contact and calculated the percentage of simulation runs with an overlapping number of *trans* interactions (i.e., 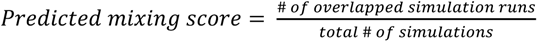). For clarification, the overlapping area was defined by the minimum number of *trans* interactions in homotypic contacts and the maximal number of *trans* interactions in heterotypic contacts. Every simulation with a number of *trans* interactions that falls within these boundaries was considered as overlapping.

## Author Contributions

G.W., N.B., and R.R designed experiments, analyzed data, and wrote the paper. G.W and K.S.A performed cell aggregation assays. G.W performed cloning, mutagenesis, western-blot, and analysis of cell aggregation assays. N.B conducted the computer simulations.

## Acknowledgments

We thank Joshua Weiner for kindly providing cPcdh plasmids. We also thank to Milana Melamed for technical help and to Barry Honig, David Sprinzak, and Nir Ben-Tal for insightful discussion. This work was supported by the Israel Science Foundation (1463/19 to R.R.).

## Supporting Information

**Fig. S1 (related to main Figure 3 and sup. Figure 2).**
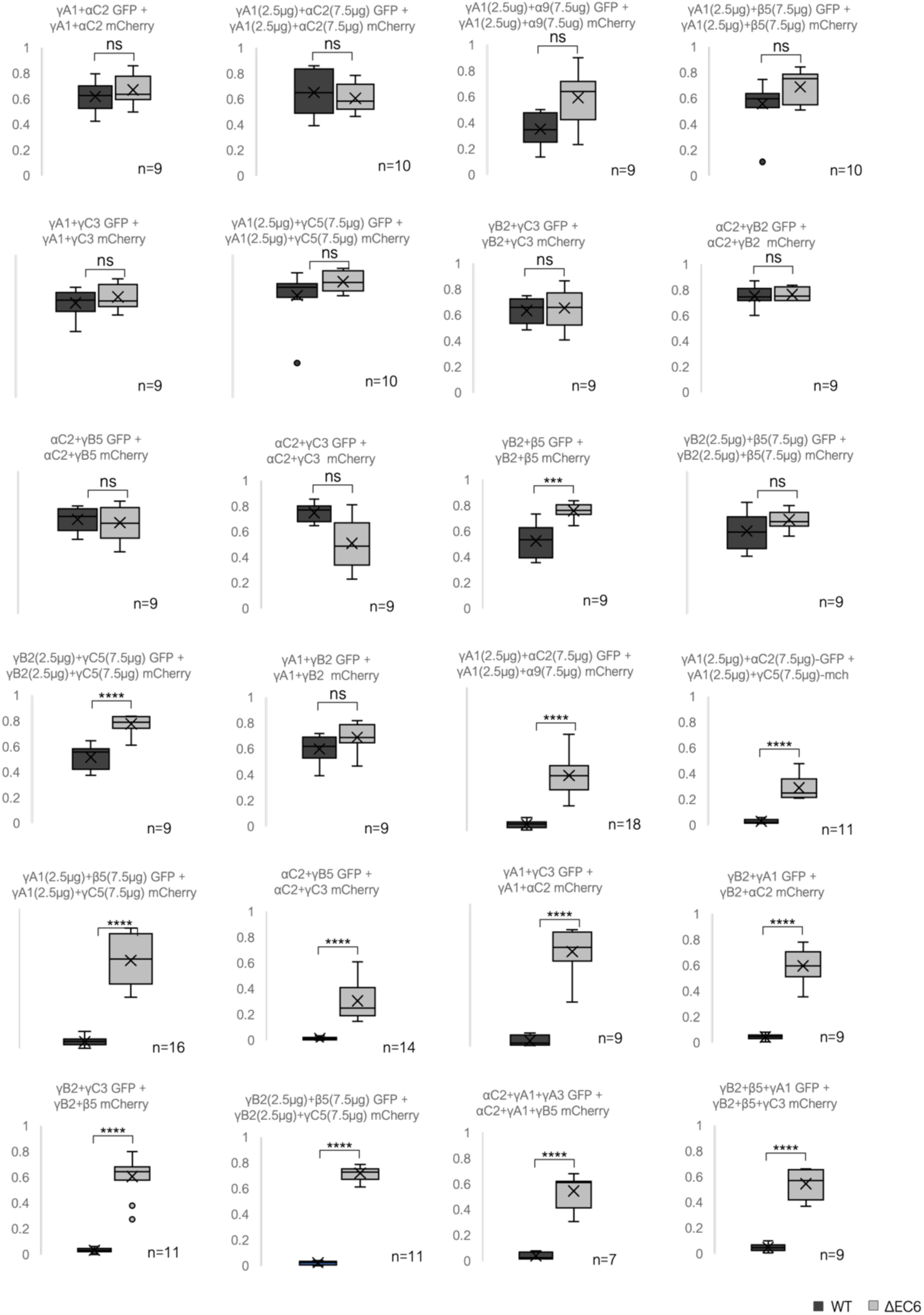
Statistical analyses of the mixing scores calculated for co-aggregation assays. Isoforms used for the cell aggregation assays are noted above the relevant plot. Mixing scores assigned in automated image analyses (see Methods) is plotted for wild-type and ΔEC6 isoforms. Significance level was calculated using a two-tailed distribution, unequal variance t-test. The notations are as follow: ns (not significant) represents p-value > 0.05; *** represents p-value < 0.001; and **** represent p-value < 0.0001.

**Fig. S2 (Related to main Figure 3).**
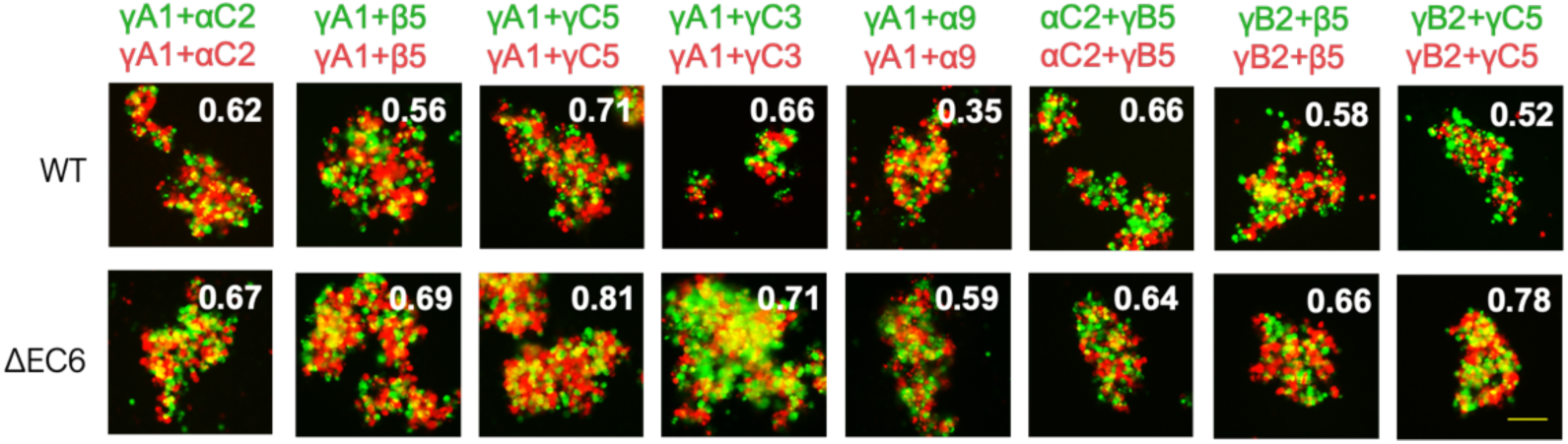
Coaggregation of pairs of matching isoforms demonstrates that eliminating *cis* interactions does not prevent *trans* interactions. (A) Results for cell aggregation assay for two K562 cell populations co-expressing pairs of matched cPcdh with wild-type (Top) or ΔEC6 ectodomains (Bottom). The mean mixing index values are shown in the upper right corner of each representative image. Cells intermixed in all cases. Scale: 100 µm

**Fig. S3 (related to Figure 6).**
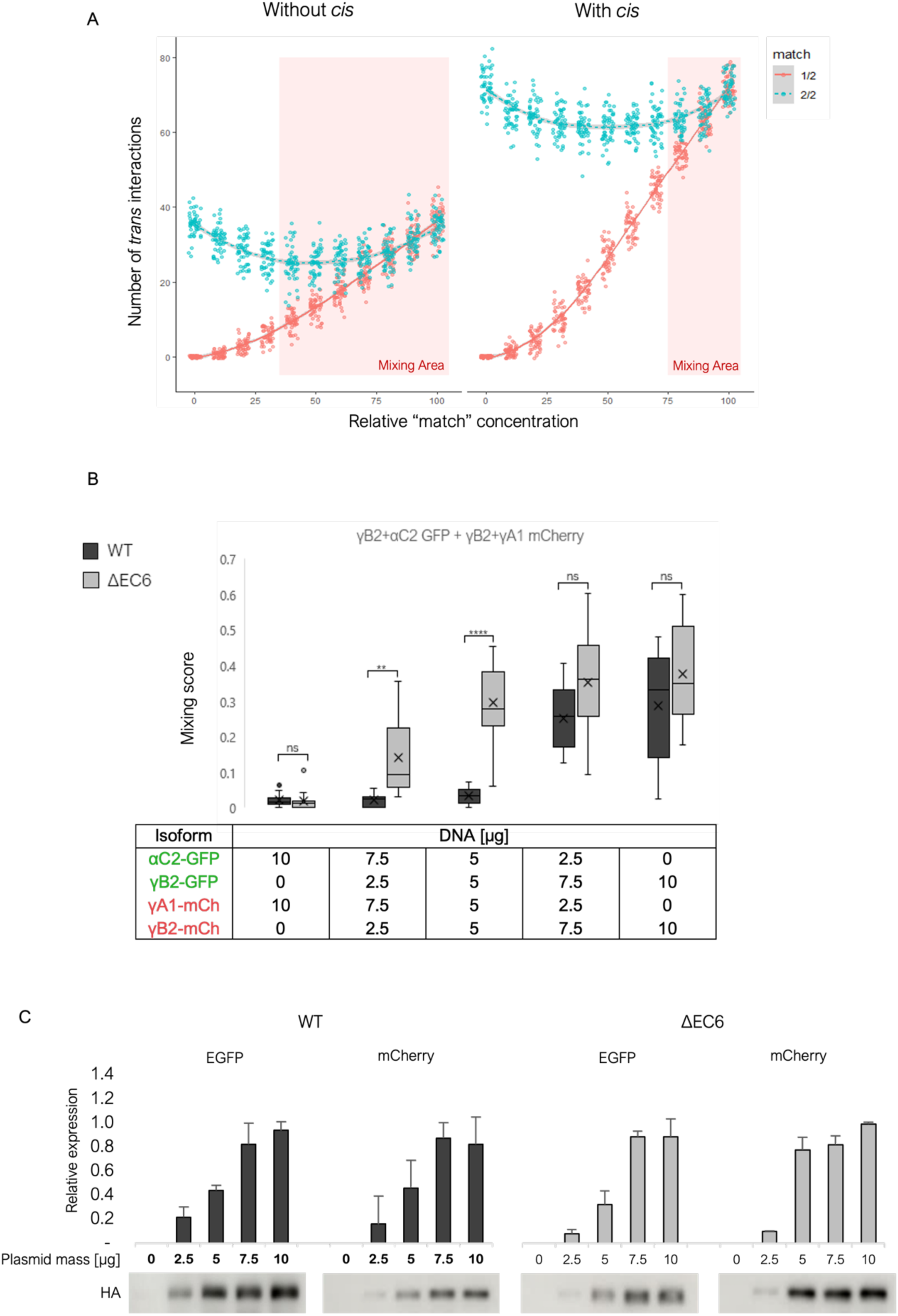
Effect of increased expression of the matching isoforms on *trans* interactions and cell population mixing. A) The change in average number of *trans* dimers as a function of increased concentrations of the matched isoforms and its dependence on *cis* interactions is plotted for all simulation runs. Blue dots represent the number of *trans* dimers for simulations between cells from the same populations and red dots represent the number of *trans* dimers for simulations between cells from different populations (containing a mismatched isoform). Highlighted in red are values of matched isoforms for which we predict mix aggregates because the number of *trans* dimers formed in the interaction of cells from the same population is similar to those formed by the interactions of cells from different populations. B) Box plot analyses of the mixing scores for triplicate cell aggregation assays shown on main Figure 6C. The amount of plasmid DNA used to transfect the K562 cells is listed in the table below. C) Western blot analyses of cPcdh expression in the cell aggregation assay shown in main Figure 6C. The “matching” isoforms, cPcdhγB2, is expressed with an HA tag.

## Notes

### Competing Interest Statement

The authors have declared no competing interest.

### Summary of Updates

correcting a technical issue with the figure resolution and image quality. No changes in data presented either in text or figures.

